# Structure of the 30S ribosomal decoding complex at ambient temperature

**DOI:** 10.1101/262972

**Authors:** E. Han Dao, Frédéric Poitevin, Raymond G. Sierra, Cornelius Gati, Yashas Rao, Halil Ibrahim Ciftci, Fulya Akşit, Alex Mcgurk, Trevor Obrinski, Paul Mgbam, Brandon Hayes, Casper De Lichtenberg, Fatima Pardo-Avila, Nicholas Corsepius, Lindsey Zhang, Matt Seaberg, Mark S. Hunter, Mengling Liang, Jason E. Koglin, Soichi Wakatsuki, Hasan Demirci

**Affiliations:** Stanford PULSE Institute, SLAC National Laboratory, Menlo Park, California, USA; Department of Structural Biology, Stanford University, Palo Alto, California, USA; Linac Coherent Light Source, SLAC National Laboratory, Menlo Park, California, USA; Biosciences Division, SLAC National Laboratory, Menlo Park, California, USA; Institutionen för Kemi, Kemiskt Biologiskt Centrum, Umeå Universitet, Umeå, Sweden

**Keywords:** Serial Femtosecond X-ray crystallography, ribosome, decoding, ambient temperature, antibiotics

## Abstract

The ribosome translates nucleotide sequences of messenger RNA to proteins through selection of cognate transfer RNA according to the genetic code. To date, structural studies of ribosomal decoding complexes yielding high-resolution data have predominantly relied on experiments performed at cryogenic temperatures. New lightsources like the X-ray free electron laser (XFEL) have enabled data collection from macromolecular crystals at ambient temperature. Here, we report an X-ray crystal structure of the *Thermus thermophilus* 30S ribosomal subunit decoding complex to 3.45 Å resolution using data obtained at ambient temperature at the Linac Coherent Light Source (LCLS). We find that this ambient-temperature structure is largely consistent with existing cryogenic-temperature crystal structures, with key residues of the decoding complex exhibiting similar conformations, including adenosine residues 1492 and 1493. Minor variations were observed, namely an alternate conformation of cytosine 1397 near the mRNA channel and the A-site. Our serial crystallography experiment illustrates the amenability of ribosomal microcrystals to routine structural studies at ambient temperature, thus overcoming a long-standing experimental limitation.

## INTRODUCTION

The bacterial ribosome possesses universally conserved functional centers that are structurally dynamic and undergo local and large-scale conformational rearrangements during protein synthesis (Ogle et al. 2001; Petrov et al. 2011; Loveland et al. 2017). In particular, the small (30S) ribosomal subunit, which is responsible for decoding the messenger RNA sequence, undergoes rearrangement of the universally-conserved monitoring residues A1492 and A1493 in the decoding center, as well as a large-scale domain closure that involves movement of the body of the whole 30S subunit closer to helix 44 (Yoshizawa et al. 1999; Ogle et al. 2001; Demirci et al. 2013a). Although such conformational changes during translation have been identified, current understanding of the structural dynamics of decoding remains incomplete. (Ogle et al. 2003; Loveland et al. 2017).

X-ray crystallography performed at synchrotron light sources has revealed the structures of the ribosome complexes at high resolution (Ban et al. 2000; Schluenzen et al. 2000; Wimberly et al. 2000; Yusupov et al. 2001; Schuwirth et al. 2005; Korostelev et al. 2006; Selmer et al. 2006; Yusupova et al. 2006; Ben-Shem et al. 2011). A key aspect to this important achievement was the development of procedures to collect data at cryogenic temperature (Haas and Rossmann 1970; Evers et al. 1994), which enabled the acquisition of datasets from single crystals that contained sufficient high-resolution data to build high-resolution models (Warkentin et al. 2014). Electron microscopy performed at cryogenic temperature has added even more information about the ribosome during several steps of protein synthesis, at increasing resolution to an existing wealth of structural information spanning a large number of species and conformational states (Mitra et al. 2005; Fischer et al. 2015; Natchiar et al. 2017). Cooling ribosomal samples to cryogenic temperature mitigates the propagation of radiation damage by reducing the movement of radicals produced by irradiation with electrons or X-rays, lowering the thermal fluctuations and conformational distributions of side- and main-chain residues, and rigidifying the structure, partly through dehydration by adding hygroscopic cryo-protectants (Warkentin et al. 2014). Therefore, while it enables successful data collection, cryo-cooling can potentially mask important details about local and global conformational dynamics and allostery of the ribosome.

The flexibility of ribosomal RNA is of significant importance in our understanding of the functional relevance of nucleic acids (Noller 2013). Additionally, the structure and conformational heterogeneity of RNA molecules are determined by the composition and physico-chemical state of the surrounding electrolytic medium (Anthony et al. 2012; Lipfert et al. 2014; Allred et al. 2017). Structural studies of ribosomes at temperatures closer to the physiological range could potentially reveal previously-obscured conformations and provide a means to evaluate their local dynamics and role in catalysis (Sierra et al. 2016).

In this work, we present an ambient-temperature structure of a 30S ribosomal decoding complex through a serial femtosecond X-ray crystallography (SFX) experiment. Using 40-femtosecond pulses, we obtain diffraction from microcrystals prior to the onset of radiation damage induced by the X-ray beam using CXI instrument at LCLS an X-ray free-electron laser (XFEL)(Liang et al. 2015). The microcrystals contained ribosomal subunits bound to a cognate mRNA-anticodon stem loop (ASL) complex and were introduced to the X-ray beam in a liquid suspension with an electrokinetic sample injector (Sierra et al. 2016). We then compared our structure to two analogous structures solved through synchrotron X-ray diffraction collected at cryogenic temperature (Ogle et al. 2001; Demirci et al. 2013a).

## RESULTS AND DISCUSSION

### SFX workflow for microcrystals of 30S ribosomal subunits

A brief outline of the experimental methods used for data collection follows, along with references to the pertinent subsections within the ***Materials & Methods***. Microcrystals of the 30S ribosomal subunit were soaked with 80 *µ*M paromomycin, 200*µ*M mRNA and 200 *µ*M phenylalanine tRNA anticodon stem loop, ASLPhe oligonucleotide, resulting in a slurry of 2 × 2 × 4 μm^3^ size 30S decoding complex microcrystals (see ***Preparation and crystallization of 30S ribosomal subunits for SFX crystallography at an XFEL***). They were delivered to the X-ray beam at the Coherent X-ray Imaging (CXI) instrument (Liang et al. 2015) of the Linac Coherent Light Source (LCLS; Menlo Park, CA, USA) via a concentric electrokinetic liquid injector (see ***coMESH construction*** and ***Operation of the coMESH*)**, as previously used for crystalline ribosome samples (Sierra et al. 2016) (Fig. 1A) (see ***Selecting a sister liquor for ribosome microcrystalline slurry*** and ***ribosome microcrystalline sample injection by using coMESH*).** During a six-hour “protein crystal screening” beamtime, we collected a complete dataset extending to 3.45 Å resolution (**Table 1**). A total of 1,731,280 detector frames were collected (corresponding to 240 minutes of net data collection time), of which 165,954 contained diffraction data. Of these frames, 19,374 patterns were indexed and merged into the final dataset. (see ***Data collection and analysis for SFX studies at LCLS*** and ***Ambient temperature 30S ribosomal subunit SFX structure refinement***).

**FIGURE 1.**
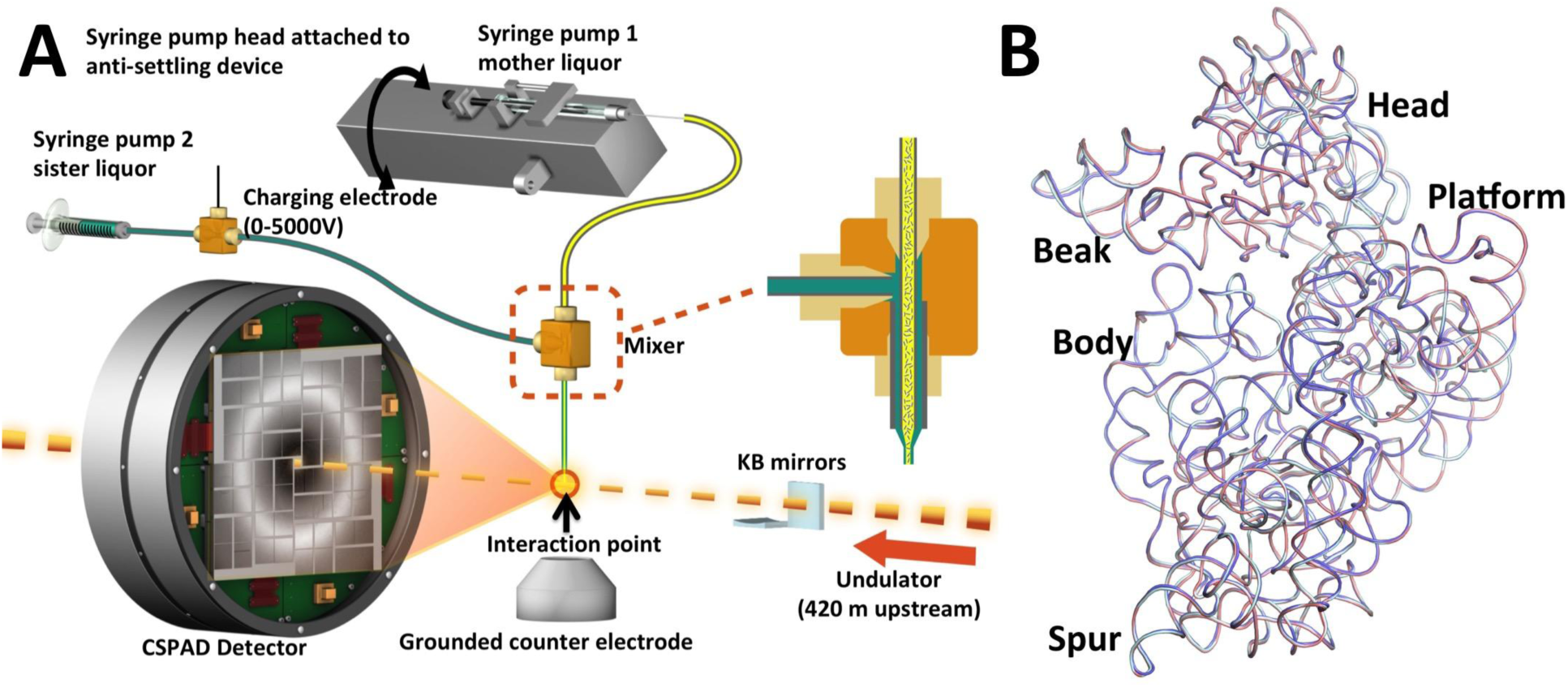
Serial Femtosecond X-ray (SFX) crystallography studies of 30S ribosomal subunit decoding complexes (A) Diagram of the concentric-flow MESH injector setup at the CXI instrument of the LCLS. The liquid jet, comprising microcrystals and their mother liquor (colored in yellow), flows in the continuous inner capillary (100 μm × 160 μm × 1.5 m; colored in gray). The sister liquor (colored in green) is charged by a high voltage power supply (0-5,000 V) for electro-focusing of the liquid jet. A mixer (indicated within the dashed orange rectangle) joins the two capillaries (colored in gray) concentrically. The sample reservoir containing ribosome microcrystals is mounted on an anti-settling device, which rotates, at an angle, about the capillary axis to keep the microcrystals suspended homogenously in the slurry. The liquid jet and the LCLS pulses interact at the point indicated by the orange circle. (B) Comparison of *T. thermophilus* 30S-ASL-mRNA-paromomycin complex structures. Superposition of 16S rRNA backbones from cryo-cooled structures colored in cyan and slate PDB IDs: 4DR4 and1IBL respectively with ambient temperature structure colored in salmon. The positions of the major 30S domains are indicated. All X-ray crystal structure figures are produced with PyMOL (http://www.schrodinger.com/pymol).

**Table 1.**
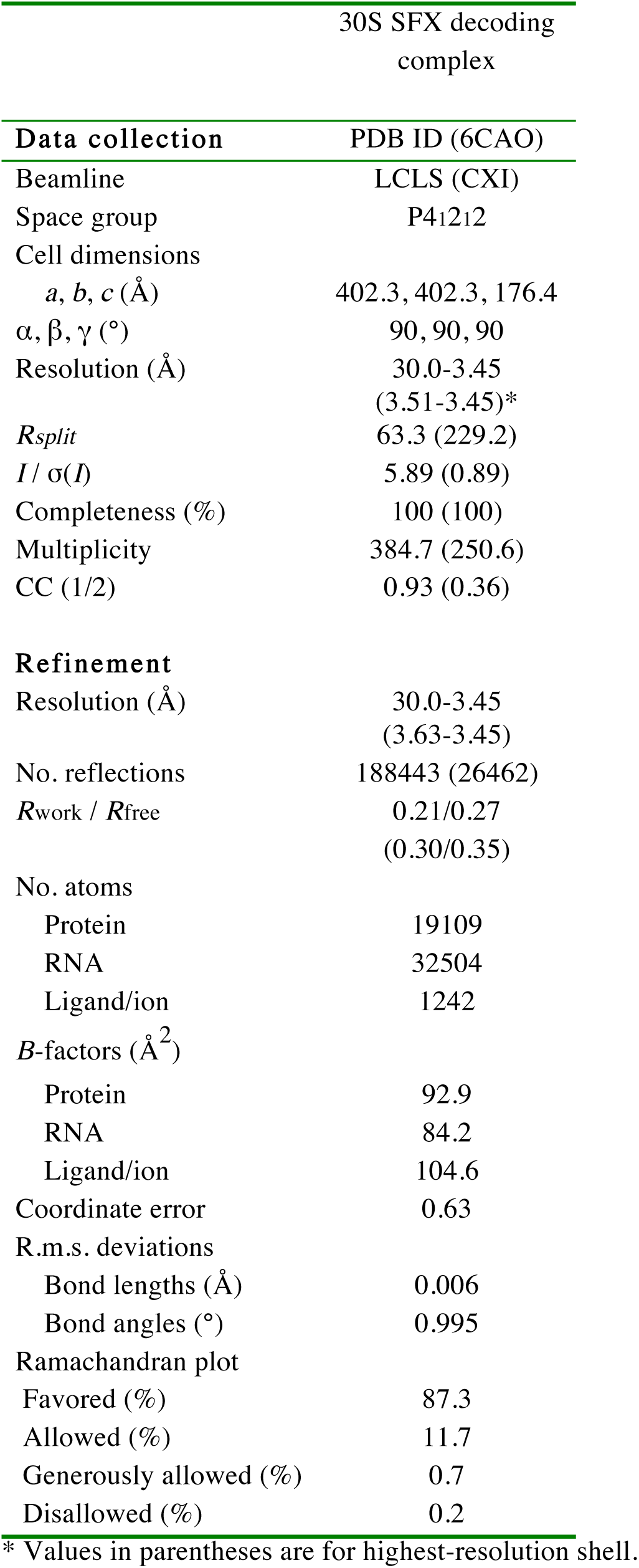
**Data collection and refinement statistics**

### Decoding complex structure at ambient temperature largely similar to prior structures

In comparing the ambient-temperature structure with its equivalents at cryogenic temperature (PDB IDs: 1IBL and 4DR4) (Ogle et al. 2001; Demirci et al. 2013a), we found that the overall decoding crystal structure was very similar to the cryogenic structures; however, we observed an alternate conformation of mRNA channel residue C1397 (further discussed below). A least-squares alignment of all 17,056 16S rRNA atoms in the 30S structures showed an overall root-mean-square deviation (RMSD) of 0.45 and 0.62 Å between the new structure and the cryogenic structures 4DR4 and 1IBL, respectively (Fig. 1B). The resulting electron density map indicated good mRNA and ASLPhe density quality in Fo-Fc difference electron density maps (Fig. 2A). The crystal structure of the 30S decoding complex at ambient temperature adopted the canonical decoding conformation, with the h44 residues A1492 and A1493 flipped out toward the minor groove of ASL and mRNA pair, consistent with the cryogenic data (Fig. 2B). The minimal binding differences observed between the cryogenic and ambient temperature 30S decoding complex structures (Figs. 1B, 2B-D) suggest that ribosomal decoding complexes can be probed at cryogenic temperature and may still be representative of what occurs at ambient temperature. However, we identified small differences within the mRNA channel in the region between the S4-S5 protein interface and the 30S acceptor A-site.

**FIGURE 2.**
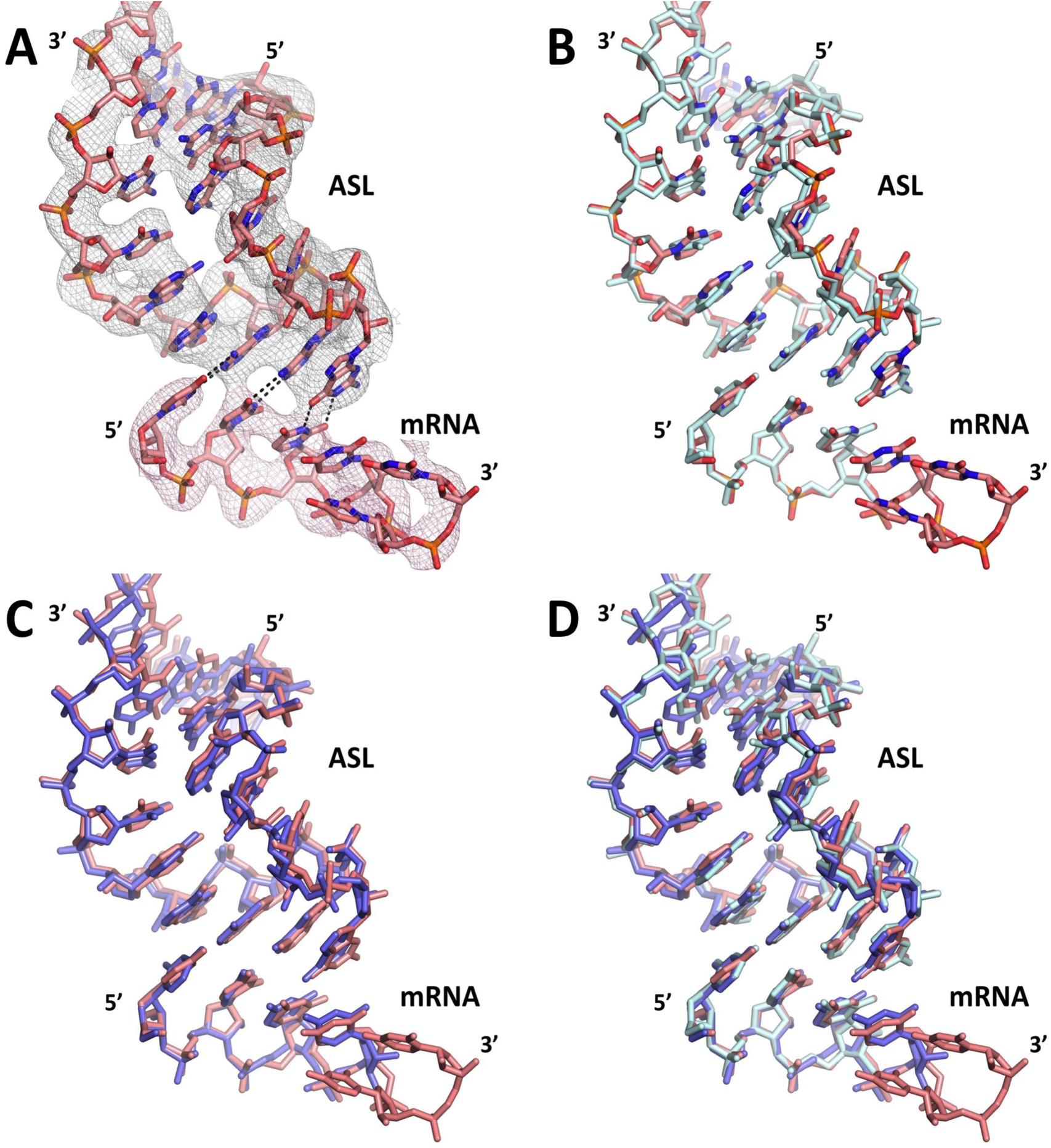
Structural comparison of ambient and cryogenic temperature decoding complexes. (A) Final unbiased Fo-Fc simple omit ambient temperature electron-density map of mRNA and ASL contoured at the 3 σ level, colored in gray and shown at 3 Å. (B) Superposition of the ambient (salmon) and cryo (cyan) temperature structures from our group showing the agreement between them. (C) Superposition of our ambient temperature structure (salmon) and the identical cryo temperature structure obtained by another laboratory (slate) (PDB ID 1IBL) showing the agreement between them. (D) Superposition of our ambient temperature structure (salmon) with the two cryo structures (cyan and slate).

### Structural perturbation in the mRNA channel

In the structure presented here, the hydrogen bonding interactions of codon residues 1-3 of the hexauridine in the mRNA-ASL complex were markedly similar to those observed in the cryogenic temperature crystal structure, as shown by the clear positive difference density for each base pair in the omit Fo-Fc electron density maps at 3.45 Å resolution (Fig. 3A-F). On the other hand, residues 4-6 of the hexa-uridine were disordered and as a result not modeled in the cryogenic structure, while a well-defined electron density was observed in the ambient-temperature structure (Fig. 2A). Furthermore, the 16S rRNA residue C1397 was captured in an alternate conformation, pointing away from the disordered hexauridine in the cryogenic dataset and involved in a stabilizing interaction with the mRNA phosphate group of residue 4 in the ambient-temperature structure (Fig. 4A-D).

**FIGURE 3.**
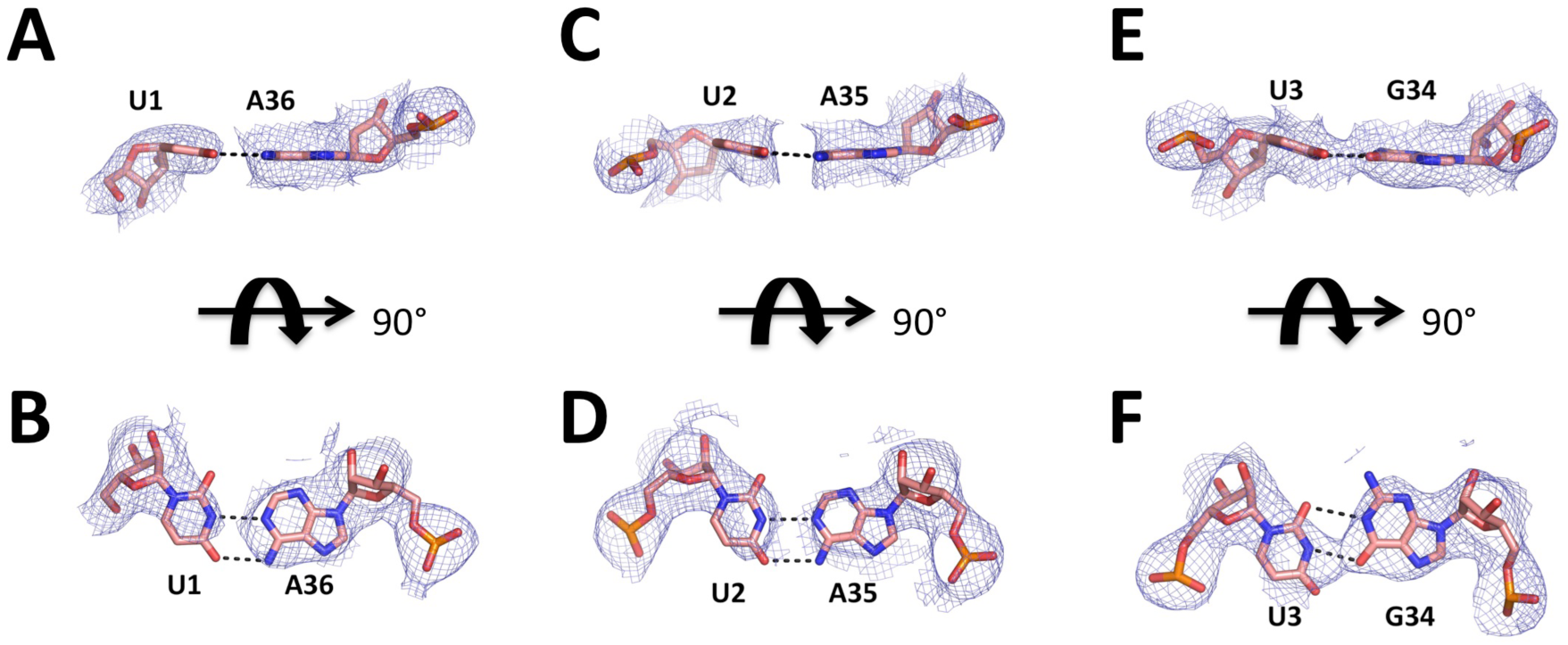
X-ray crystallography structures of the decoding mRNA-ASL minihelix. (A) Simple Fo-Fc omit maps of the decoding complex individual base pair mRNA(U1)-ASL(A36) contoured at the 3 σ level, colored in gray and shown at 3 Å. (B) same as A rotated around x-axis by 90 degrees. (C) Simple Fo-Fc omit maps of the decoding complex individual base pair mRNA(U2)-ASL(A35) contoured at the 3 σ level, colored in gray and shown at 3 Å. (D) same as C rotated around x-axis by 90 degrees. (E) Simple Fo-Fc omit maps of the decoding complex individual base pair mRNA(U3)-ASL(G34) contoured at the 3 σ level, colored in gray and shown at 3 Å. (E) same as C rotated around x-axis by 90 degrees.

**FIGURE 4.**
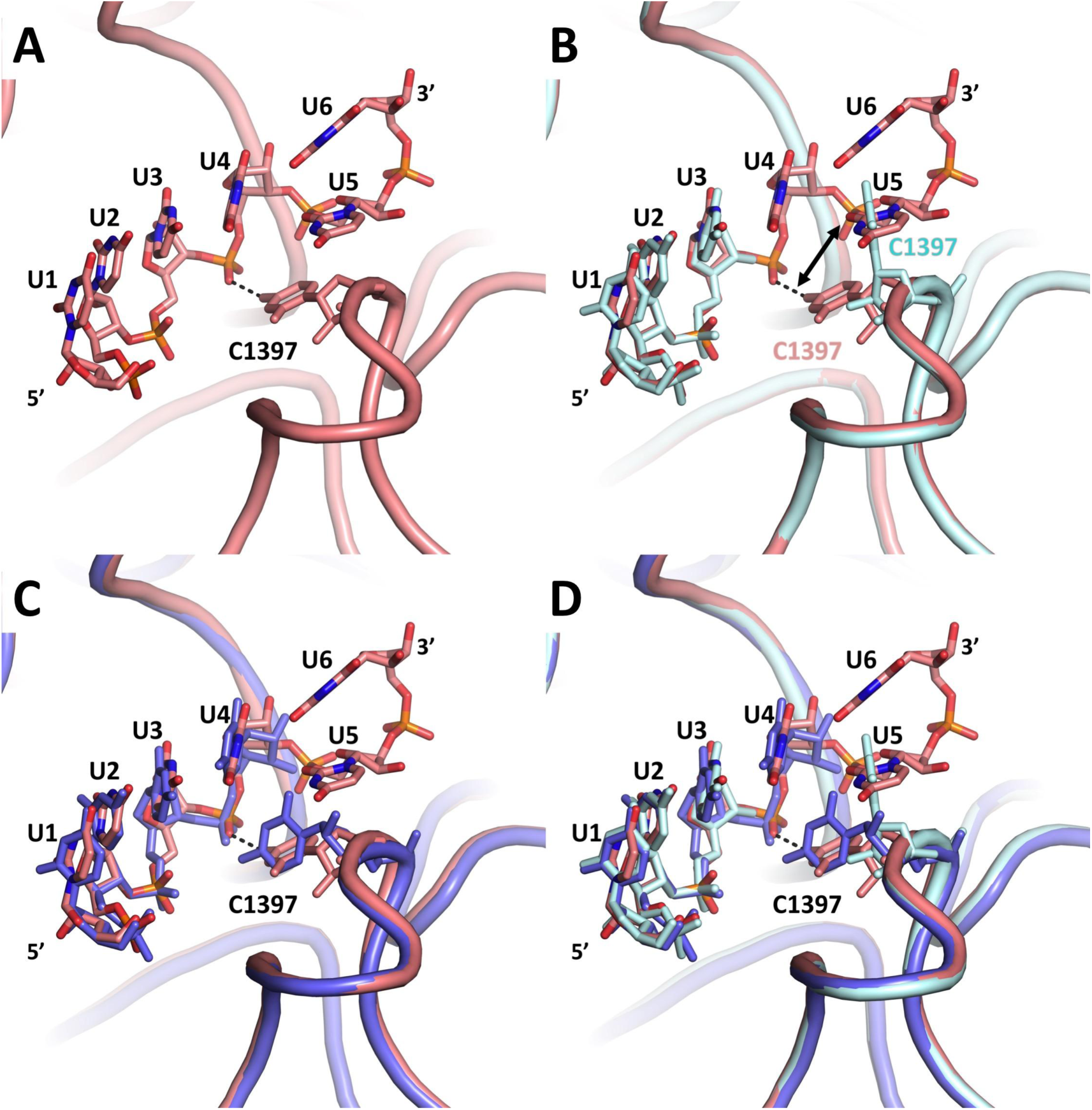
Temperature dependence of C1397 conformational dynamics. (A) Ambient temperature mRNA model at the decoding region. C1397 engages in H-bonding with U4 which demarcates the boundary between the A-site mRNA codon and codon −1. (B) Superposition of the ambient temperature 30S-decoding complex with cryogenic structure shows the disorder in the cryogenic mRNA structure also the alternate conformation of the C1397. (C) Superposition of our ambient temperature structure (salmon) and the identical cryo temperature structure obtained by another laboratory (slate) (PDB ID 1IBL) showing the disorder in the cryogenic structure. (D) Superposition of our ambient temperature structure (salmon) with the two cryo structures (cyan and slate).

The movement of the messenger RNA is controlled during the transition from pre- to post-translocation (Moazed and Noller 1989; Achenbach and Nierhaus 2015). The universally conserved bases C1397 and A1503 of the 16S rRNA head domain located on top of large secondary rRNA structures (Zhou et al. 2013; Achenbach and Nierhaus 2015) and respectively intercalate between nucleotide pairs +9 and +10 and −1 and −2 of the mRNA, exclusively in the intermediate states of translocation. Those two residues are thought to prevent a back-sliding of the mRNA during back-rotation of the 30S head, thus exerting a pawl function. In the post-translation state, however, the two intercalating nucleotides do not touch the mRNA. Residue C1397 has been described as being able to adopt multiple conformations in several crystal structures, in response to the presence of a tRNA at the A-site (Jenner et al. 2010) (Zhou et al. 2013). The comparison of our ambient and cryogenic temperature structures therefore illustrates the intrinsic flexible nature of C1397 even in the presence of tRNA at the A-site.

### Ambient-temperature studies build upon previous cryo-crystallography data

The low RMS deviation between the crystal structures collected in this work and its equivalents studied at cryogenic temperature suggests that previous structural data have captured the 30S ribosomal subunit in a representative conformation, although the causes of the observed discrepancies between datasets are not immediately apparent. Our demonstration of data collection at ambient temperature indicates that the long-standing limitation of relying on cryogenic temperatures to study ribosome decoding complexes at high resolution is surmountable with contemporary approaches. This latest effort builds upon prior attempts by our group to obtain a full dataset at an XFEL. One factor we felt to be important to the success in data collection here was the use of an electrokinetic injector, the “concentric MESH” (Sierra et al. 2016). This sample injector delivers microcrystals to the X-ray beam by relying on electric charge rather than pressure and allowed us to simplify the apparatus as described (Figure 1A). The time needed for data collection, four hours, suggests that a full dataset can be collected within a beamtime shift at any XFEL. With additional improvements in real-time data processing and the use of high-repetition-rate sources such as European XFEL, the amount of time needed could potentially be further reduced.

A major promise of structural studies performed at ambient temperature is the ability to probe biological macromolecules at near-physiological temperatures in liquid suspension. Time-resolved SFX is an emerging approach that builds on the successes of SFX in determining static structures and has been demonstrated for photo-sensitive targets (Tenboer et al. 2014; Pande et al. 2016), with a time resolution as short as 100 fs. Experimental approaches to mix enzymes with their substrates on relevant enzymatic time scales in real-time prior to their delivery to the X-ray beam are also under development and would enable the probing of structural intermediates of these enzymes (Kupitz et al. 2017). The first demonstration of time-resolved structural enzymology at an XFEL has been performed on a riboswitch and its substrate with a mixing time of 10 second (Stagno et al. 2017), paving the way to performing similar experiments on ribonucleo-protein complexes such as the ribosome. Time-resolved studies of the ribosome decoding complexes could potentially elucidate the process of decoding to a level of clarity not previously attainable and answer key questions about the structural basis of decoding dynamics.

## MATERIALS AND METHODS

### Preparation and Crystallization of 30S Ribosomal Subunits for SFX crystallography at an XFEL

30S ribosomal subunits from *T. thermophilus* HB8 (ATCC27634)(Yoshizaki et al. 1971) were prepared as previously described (Wimberly et al. 2000; Demirci et al. 2010). Purified 30S ribosomal subunits were crystallized at 4 °C by the hanging drop method using a mother liquor solution containing 17% (v/v) 2-methyl-2,4-pentanediol (MPD), 15 mM magnesium acetate, 200 mM potassium acetate, 75 mM ammonium acetate and 100 mM 2-(*N*-morpholino) ethanesulfonic acid (MES)-KOH (pH 6.5). The mRNA fragment (with codon sequences underlined), 5‘<u>UUU</u>UUU3’ and the anticodon stem loop ASLPhe, (with anticodon sequence underlined GGGGAUU<u>GAA</u>AAUCCCC, were purchased from Dharmacon (GE Life Sciences). Microcrystals 2 × 2 × 4 μm^3^ in size were harvested in the same mother liquor composition, pooled (total volume of 500 μl loaded) and supplemented with 200μM of mRNA oligomer, ASLPhe and 80 μM paromomycin for 48 hours before data collection. The crystal concentration was approximated to be 10^10^-10^11^ particlesper ml based on light microscopy and nanoparticle tracking analysis (NanoSight LM10-HS with corresponding Nanoparticle Tracking Analysis (NTA) software suite (Malvern Instruments, Malvern, UK).

### coMESH construction

The concentric microfluidic electrokinetic sample holder (coMESH), which is the liquid injector used to deliver the microcrystals to the X-ray beam, is comprised of two capillaries: one large and one small. The central, smaller capillary carrying the microcystals of the 30S decoding complex was a continuous fused silica capillary of 100 *µ*M inner diameter, 160 *µ*M outer diameter, and 1.5 m length (Polymicro, Phoenix, AZ, USA). This capillary directly connected the sample reservoir with the X-ray interaction region, passing through a vacuum feedthrough and then through a microfluidic *Interconnect Cross* (C360-204, LabSmith, Livermore, CA USA; C360-203 *Interconnect Tee* may also be used with no need to plug the fourth channel) (Fig. 1A). The uninterrupted, straight capillary was free of in-line filters, unions or any other connections, thus minimizing the risk of sample clogs and leaks, which most often occur at flow channel transitions. A similar uninterrupted approach was used for the larger capillary. The fitting at the top of the *cross* in Figure 1A was an adapter fitting (T116-A360 as well as T116-100, LabSmith, Livermore, CA, USA) which compressed a 1/16” (1.6 mm) OD, 0.007” (178 *µ*M) ID polymer (FEP) tubing sleeve (F-238, Upchurch, Oak Harbor, WA, USA) onto the sample capillary. This connection method was necessary to properly seal the capillary to the *cross*. The sheath line at the bottom of the figure was a short 5 cm fused silica capillary with an outer diameter of 360 μm and an inner diameter of 180, 200, or 250 *µ*M depending on the desired flow rate of the sheath liquid at a given driving pressure. This outer capillary was connected to the *cross*, and the sheath liquid was supplied from the third port of the cross (the left port in the figure) through a compatible microfluidic tubing; we typically used silica capillaries with 360 *µ*M outer diameters and 200 *µ*M inner diameters for optimum flow rate. For simplicity, the injector has been designed to operate at relatively low backing pressures (up to a few atmospheres). For this experiment, the sheath liquid line was connected to a syringe filled with the appropriate sister liquor which contains the cryoprotection solution, driven by a syringe pump (PHD Ultra, 703006, Harvard Apparatus, Holliston, MA, USA) We electrically charged the sheath liquid between 0-5,000 V potential using an in-line conductive charging union (M-572, Upchurch, Oak Harbor, WA, USA) connected to a SRS PS350 (Stanford Research Systems, Sunnyvale, CA, USA) high voltage source.

The capillary assembly was loaded into a vacuum chamber with 1 × 1 *µ*M^2^ focused XFEL beam and the standard load-lock system of the CXI instrument. A grounded, conical counter electrode with a 1 cm opening was placed approximately 5 mm below the capillary tip; the capillaries and opening of the cone were coaxial. The angle of the cone and its distance from the tip were set to enable a diffraction cone with a 45° 2q scattering angle. All capillaries were fed through vacuum flanges with 1/16” (1.6 mm) Swagelok bulkhead fittings using appropriately sized polymer sleeves. The sheath reservoir was a 1 ml Gastight Hamilton syringe with a PTFE Luer tip (1001 TLL SYR, Hamilton, USA). The ribosome microcrystalline sample was supplied from a 500 *µ*l Hamilton Gastight syringe.

### Selecting a sister liquor for the ribosome microcrystalline slurry

The coMESH incorporates a “sister liquor” as part of its sample delivery design which is used to protect the microcrystals and their mother liquor from the dehydrative effects of the sample chamber vacuum (10^−5^ Torr). This fluid travels in the larger capillary and, after reaching the cross, peripherally to the sample line until the terminus of the capillary within the sample chamber. The choice of fluid may reflect the composition of the crystals’ mother liquor but need not strictly match it. For delivery of ribosome microcrystals using co-terminal capillaries (Sierra et al. 2016), the compatibility between the sister liquor and the ribosome crystals was not considered since fluid interaction occurred immediately prior to reaching the X-ray interaction region, allowing minimal time, if any, for the fluids to mix. Current and prior experimental experience suggest that a 26% MPD-containing sister liquor seems to be an ideal choice for co-terminal sample delivery. The buffer has proven to be quite reliable, surviving injection into vacuum with minimal issues especially with respect to buildup of dehydrated precipitates, which can occur with salts or polyethylene glycols and is problematic for sample delivery due to its adherence to (and thus, blockage of) the capillary tip. Other factors influencing the choice of fluid include availability of the fluid and its viscosity. The lower viscosity allows the sister liquor to be pumped easily by a standard syringe pump through the more resistive *cross* manifold or concentric annular flow before the exit region of the capillary.

### Operation of the coMESH

Before connecting the central sample line, the sister liquor was loaded, flowed and electrically focused. Once a slightly stable jet was achieved, the central sample line carrying ribosome slurry was connected. Notably, the sister liquor never fully stabilized because of the entrained air from the disconnected sample line continuously introducing bubbles. The central sample line had much less fluidic resistance compared to the outer line; connecting it first with a vacuum sensitive sample will cause immediate jet clogging and blockages, for this reason the outer line should always be on while operating in vacuum environment. In general, the sister liquor was set to flow at or near the flow rate of the mother liquor. If diffraction hits were not observed, the sister liquor flow was reduced and/or the mother liquor flow was increased slowly (to ensure stable jetting) until diffraction patterns reappeared.

### Ribosome microcrystalline sample injection with coMESH

For the study of 30S-mRNA-ASL-paromomycin complex crystals, the inner sample line contained unfiltered crystals in their native mother liquor containing 17% (v/v) 2-methyl-2,4-pentanediol (MPD). The size distribution of the 30S crystals was uniform owing to their controlled slower growth at 4°C (Demirci et al. 2013b). Occasional larger-sized 30S crystals were discarded by repeated gentle differential settling without centrifugation. The outer sister liquor was the same buffer, with no crystals, as the mother liquor in that the original substituent concentrations remained constant while having increased the MPD to 26% (v/v) to aid in-vacuum injection.

### Data collection and analysis for SFX studies at LCLS

The serial femtosecond X-ray crystallography (SFX) experiment was performed at Coherent X-ray Imaging instrument (CXI) of the Linac Coherent Light Source (LCLS; SLAC National Accelerator Laboratory, Menlo Park, CA, USA) under beamtime ID: cxil1416. The LCLS X-ray beam with a pulse duration of approximately 50 fs was focused using X-ray optics in a Kirkpatrick-Baez geometry to a beam size of 1.3 × 1.3 μm^2^ full width at half maximum (FWHM) at a pulse energy of 2.9 mJ, a photon energy of 9.5 keV (1.29 Å) and a repetition rate of 120 Hz.

A total of 1,731,280 images were collected with the Cornell-SLAC Pixel Array Detector (CSPAD) (Pietrini and Nettelblad 2017) from crystals of the 30S-mRNA-ASL-paromomycin complex. The total beamtime used for this dataset was 240 minutes (out of an available 360 minutes). Crystal hits were defined as frames containing more than 30 Bragg peaks with a minimum signal-to-noise ratio greater than 4.5 on each detector image, for a total of 165,954 hits. The detector distance was set to 223 mm, with an achievable resolution of 3.08 Å at the edge of the detector (2.6 Å in the corner).

After the detection of crystal hits and following the conversion of individual diffraction patterns to the HDF5 format by *CHEETAH* (Barty et al. 2014), the *CrystFEL* software suite (White et al. 2016) was used for crystallographic analysis. The information of peak positions was used for the indexing of individual, randomly oriented crystal diffraction patterns using FFT-based indexing approaches. The detector geometry was refined using an automated algorithm to match found and predicted peaks to subpixel accuracy (Yefanov et al. 2015). The integration step was performed using a built-in Monte-Carlo algorithm to estimate accurate structure factors from thousands of individually measured Bragg peaks (Kirian et al. 2010). After the application of per-pattern resolution cutoff, frames which did not match to an initial merged dataset with a Pearson correlation coefficient of less than 0.4, were excluded from the final dataset. Diffraction intensities from a total of 19,374 indexed diffraction patterns (11.8 % indexing rate) were merged into a final dataset (Overall CC* = 0.93; 3.45 Å cutoff) for further analysis (P4_1_2_1_2, unit cell: a = b = 402.3 Å, c = 176.4 Å; α = β = γ = 90°). The final resolution cutoff was estimated to be 3.45 Å using a combination of *CC1/2* (Karplus and Diederichs 2012) and other refinement parameters. The final dataset had an overall *R*_*split*_ = 63.3% and *CC\** = 0.36 in the highest resolution shell.

### Ambient temperature 30S Ribosomal Subunit SFX Structure refinement

We determined the ambient temperature structure of the 30S-mRNA-ASL-paromomycin complex using the automated molecular replacement program *PHASER* (McCoy et al. 2007) with the previously published cryo-cooled 30S ribosomal subunit-mRNA-ASL-paromomycin complex synchrotron structure as a search model (PDB entries 4DR4 and 1IBL)(Ogle et al. 2001; Demirci et al. 2013a). The resulting structure was refined with rRNA modifications, which are mostly located near the decoding center. This choice of starting search model minimized experimental variations between the two structures such as sample preparation, crystal growth, data processing, and model registry. Coordinates of the 4DR4 with additional RNA and protein modifications were used for the initial rigid body refinement with the *PHENIX* software package (Adams et al. 2010). During the initial refinement of ambient temperature XFEL structure, the entire mRNA and ASL components were omitted and the new models were built into unbiased difference density. After simulated-annealing refinement, individual coordinates, three group B factors per residue, and TLS parameters were refined. Potential positions of magnesium or potassium ions were compared with those in the high-resolution (2.5 Å) 30S subunit structure (PDB accession code 2VQE)7 in program *COOT* (Emsley and Cowtan 2004), and positions with strong difference density were retained. All magnesium atoms were replaced with magnesium hexahydrate. Water molecules located outside of significant electron density were manually removed. The Ramachandran statistics for this dataset (most favored / additionally allowed / generously allowed / disallowed) are 87.3 / 11.7 / 0.7 / 0.2 % respectively. The structure refinement statistics are summarized in Table 1. Structure alignments were performed using the alignment algorithm of *PyMOL* (www.schrodinger.com/pymol) with the default 2σ rejection criterion and five iterative alignment cycles. All X-ray crystal structure figures were produced with *PyMOL.*

## DATA DEPOSITION

The coordinates have been deposited in the Protein Data Bank with the accession code 6CAO.

## ACKNOWLEDGEMENTS

The authors thank Michelle Young for her support and critical reading of the manuscript. Portions of this research were carried out at the Linac Coherent Light Source (LCLS) at the SLAC National Accelerator Laboratory. Use of the LCLS is supported by the U.S. Department of Energy (DOE), Office of Science, Office of Basic Energy Sciences (OBES) under Contract No. DE-AC02-76SF00515. The LCLS is acknowledged for beam time access under experiment no. cxil1416. EHD, RGS and HD acknowledge the support of OBES through the AMOS program within the CSGB and the DOE through the SLAC Laboratory Directed Research and Development Program. EHD acknowledges financial support from the Stanford University Dean of Research. HD acknowledges support from NSF Science and Technology Centers grant NSF-1231306 (Biology with X-ray Lasers, BioXFEL). FP, NC and FPA are supported by National Institutes of Health (NIH) grant R35GM122543. Parts of the sample injector used at LCLS for this research was funded by the National Institutes of Health, P41GM103393, formerly P41RR001209. CG acknowledges SLAC and the DOE for financial support through the Panofsky fellowship. CDL acknowledges Stiftelsen JC Kempes Minnes Stipendiefond for financial support. We thank A. Fry’s continuous support for LCLS intern students who contributed to the portions of the experiments carried at LCLS. H.D. acknowledges valuable discussions with A. Takeuchi, K. Dursuncan and E. Satunaz. We thank G. Stewart of SLAC for excellent technical assistance with creating graphics for Figure 1A.

## Author contributions

**HD** and EHD designed the project. HD, EHD, YR, HC, FA prepared the samples for SFX studies. RGS, HD, AM, PM, BH, TO, CDL and EHD build the coMESH injectors. FP and CG executed the SFX data reduction. NC and FPA provided realtime XFEL data analysis. HD refined the structure. The experiments were executed by HD, EHD, RGS, FP, NC, FPA, MH, ML, MS, FA, HIC, LZ, YR. Data were analyzed by HD, EHD, FP, SW, CG and RGS. The manuscript was prepared by HD, EHD, FP, CG, SW and RGS with input from all the coauthors.

